# Auditory accessory stimulus boosts pupil-linked arousal and reduces choice bias

**DOI:** 10.1101/2022.08.28.505585

**Authors:** B.M. Bruel, V.G. Katopodis, R. de Vries, T.H. Donner, M.J. McGinley, J.W. de Gee

## Abstract

Recent work indicates that pupil-linked phasic arousal signals reduce the impact of prior expectations and biases on decision formation. It has remained unclear whether phasic arousal (i) causes the bias reduction, if (ii) choosing against one’s bias causes phasic arousal, or if (iii) a third variable is driving both. Here, using an auditory accessory stimulus, we found evidence for the first scenario: on accessory stimulus vs normal trials, pupil-linked phasic arousal was robustly elevated and choice bias was reduced. With computational modeling of behavior, we established that the bias reduction was not due to a change in response caution (i.e., speed-accuracy tradeoff), but due to a change in a bias in the accumulation of evidence leading up to a choice. Thus, pupil-linked phasic arousal shapes choice behavior.

## Introduction

Most decisions are based on the accumulation of decision-relevant evidence combined with a bias (prior belief) (Shadlen & Kiani, 2013; Summerfield & de Lange, 2014): e.g., you will likely choose to buy apples when they look good and are on sale (discrete pieces of evidence) and you happen to like apples (bias). These decision-computations are implemented in a distributed choice-network of brain regions (Steinmetz et al., 2019; van Vugt et al., 2018): during judgments about weak sensory signals in noise, sensory cortex encodes the noise-corrupted decision evidence, and downstream regions of association and motor cortices accumulate this noisy sensory signal into a decision variable that determines the behavioral choice (Bogacz et al., 2006; Shadlen & Kiani, 2013; Siegel et al., 2011; Wang, 2008); these same areas control bias through fluctuations in ongoing (pre-stimulus) activity (de Lange et al., 2013; Mochol et al., 2021; van Vugt et al., 2018) and/or through asymmetries in the rate in which stimulus information is accumulated (Hanks et al., 2011; Jun et al., 2021; Kloosterman et al., 2019; Mochol et al., 2021).

Subcortical neuromodulatory systems like the noradrenergic locus coeruleus and the cholinergic basal forebrain respond briefly every time we make a decision (Aston-Jones & Cohen, 2005; Breton-Provencher et al., 2022; de Gee et al., 2017; Gritton et al., 2016), and can shape information processing in the brain’s choice network in profound ways (Aston-Jones & Cohen, 2005; Froemke, 2015; Harris & Thiele, 2011; Lee & Dan, 2012; McCormick et al., 2020). Precisely *how* these brief neuromodulatory signals shape decision computations and choice behavior is a challenging question, due to the difficultly of assaying and manipulating subcortical neuromodulatory nuclei, especially in awake animals or non-invasively in humans.

Pupillometry is a powerful tool, as fluctuations in pupil size at constant luminance track noradrenergic and cholinergic activity (Breton-Provencher & Sur, 2019; de Gee et al., 2017; Joshi et al., 2016; Mridha et al., 2021; Murphy et al., 2014; Reimer et al., 2016; Varazzani et al., 2015). The pupil dilates every time we engage in a decision (Cheadle et al., 2014; de Gee et al., 2014; Gilzenrat et al., 2010) and predicts a specific change in decision-making based on protracted evidence accumulation: a linear reduction of choice bias (de Gee et al., 2017, 2020; Lewandowska et al., 2019; Schriver et al., 2020). This relationship between task-evoked pupil responses and choice bias generalizes across species (humans and mice), signs of bias (liberal and conservative) and domains of decision-making (based on visual, auditory or memory information) (de Gee et al., 2020). Related, in dynamic environments, evoked pupil responses are associated to violations of learned, top-down expectations (Filipowicz et al., 2020; Krishnamurthy et al., 2017; Murphy et al., 2021; Nassar et al., 2012). An important limitation of all these studies is their correlational nature (but see (Nassar et al., 2012)). Thus, it remains unclear whether phasic arousal causes the bias reduction, if choosing against one’s bias causes phasic arousal, or if a third variable is driving both.

One promising way to experimentally manipulate pupil-linked arousal and choice behavior is using an accessory stimulus. It has been known for decades that humans respond faster to a visual stimulus when it is immediately preceded by a task-irrelevant auditory accessory stimulus, often without an associated change in accuracy (Bernstein et al., 1969; Hackley et al., 2009; Hackley & Valle-Inclán, 1998, 1999; Hershenson, 1962; Jepma et al., 2009; Lippert et al., 2007; Miller et al., 1999; Nickerson, 1973; Stahl & Rammsayer, 2005; Stoffels et al., 1985; Tona et al., 2016). The accessory stimulus effect (and related phasic alerting effects) might be mediated by phasic locus coeruleus activity and the subsequent release of noradrenaline in its target structures (Hackley & Valle-Inclán, 1999; Joshi et al., 2016; Joshi & Gold, 2022). In line with this, accessory stimuli boost pupil-linked arousal during decisions (Petersen et al., 2017; Tona et al., 2016). However, the effect of accessory stimuli on choice bias has not been systematically investigated (but see (Lovelace et al., 2003)).

Here, we tested the hypothesis that an accessory stimuli boosts phasic pupil-linked arousal and reduce the impact of biases on choice behavior. Subjects performed an elementary contrast detection task and heard a task-irrelevant auditory accessory stimulus (white noise) on a random one-sixth of trials. Pupil-linked arousal was consistently boosted on the accessory stimulus compared to normal trials, and choice bias was reduced. Computational modeling of behavior showed that this effect was due to a change in the bias that existed in the evidence accumulation process itself, and not in its starting point. We conclude that phasic pupil-linked arousal causally reduces the impact of biases on choice behavior.

## Results

### Accessory stimulus reduces reaction time, without an associated change in accuracy

Subjects performed a visual detection task, which consisted of a baseline interval, a subsequent decision interval and an inter-trial-interval of variable duration (**Fig. 1A**; Methods). Low-contrast dynamic random noise was continuously present throughout the decision interval. On half of the trials, a low contrast vertical grating, the signal, was superimposed onto the noise during the decision interval (**Fig. 1A**). Subjects had to report a choice about the presence or absence of the signal (yes or no) by pressing one of two buttons. During a pseudo-random one-sixth of trials, auditory white noise was presented at 70 dB from the start of the baseline interval until the end of the decision interval, which was terminated by the subject’s choice (**Fig. 1A**). To prevent the auditory accessory stimulus from acting as a temporal cue, on all trials the color of the fixation dot changed from grey to green at the start of the baseline interval (**Fig. 1A**).

**Figure 1.**
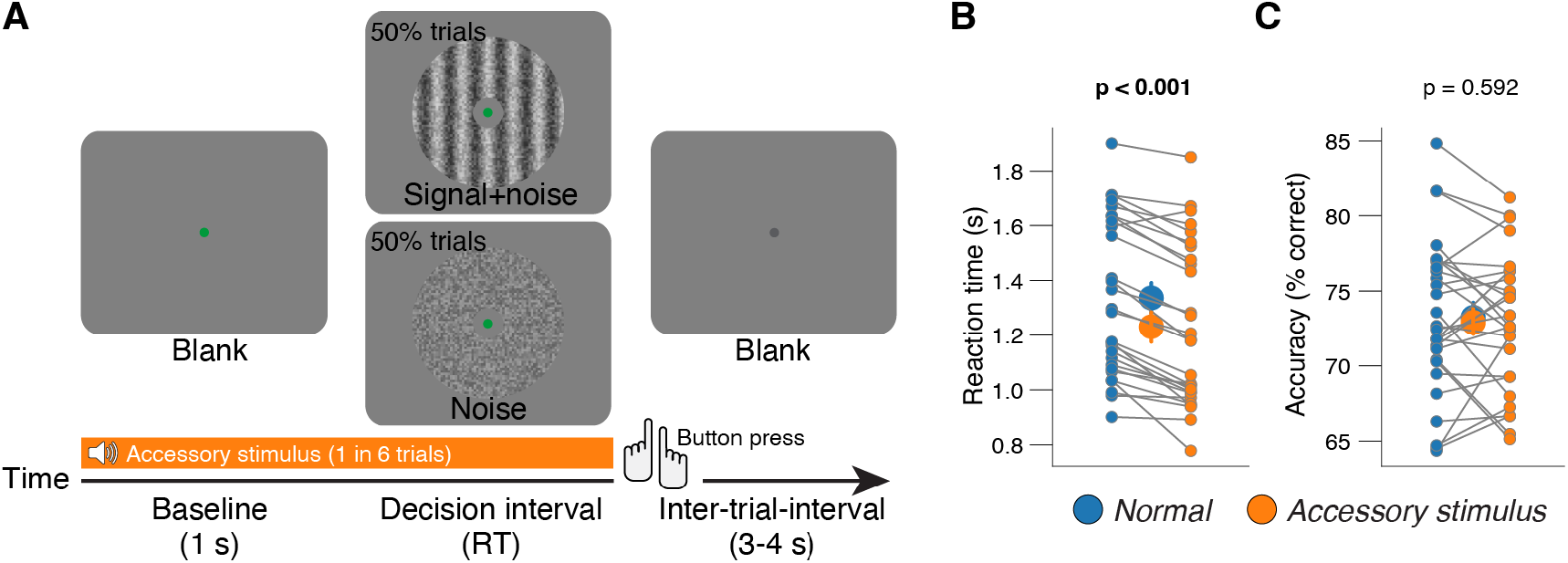
Behavioral task and task-evoked pupil responses. **(A)** Schematic sequence of events during the yes-no contrast detection task. Subjects reported the presence or absence of a faint grating signal superimposed onto dynamic noise. Signal contrast is high for illustration only. **(B)** Reaction time, separately for normal and accessory stimulus trials. Every connecting line is a subject; large data points in the middle are the group averages. Error bars, 68% confidence interval across subjects (N=29); stats, paired sampled t-test. **(C)** As B, but for accuracy.

The signal contrast was adjusted individually such that each subject performed at roughly 75% correct on the normal trials (**Fig. 1B**). When pooling normal trials across all participants, the median reaction time was 1.26 s (range across individuals: 0.83-1.87 s) (**Fig. 1C**). Thus, subjects produced long reaction times consistent with a protracted decision process.

We first sought to establish the typical accessory stimulus effect: a reduction of reaction time, without an associated change in accuracy. In line with this, on accessory stimulus vs normal trials, reaction times were shorter in all subjects, except for one (**Fig. 1B**). Median reaction times sped up by an average (± S.E.M.) of 105 ms (13 ms). There was no consistent difference in accuracy (**Fig. 1C**).

In sum, our visual detection task produced protracted decisions and the typical accessory stimulus effects on choice behavior.

### Accessory stimulus boosts pupil-linked arousal throughout decision formation

During perceptual decisions, pupil-linked arousal is transiently elevated (de Gee et al., 2014). Trial-to-trial fluctuations in the magnitude of such phasic pupil responses are primarily due to the difference in a sustained component that spans the entire interval from decision interval onset to behavioral choice (de Gee et al., 2017). Therefore, with an auditory accessory stimulus we aimed to boost pupil-linked arousal throughout the protracted decision process.

Pupil size fluctuated at a relatively slow time scale throughout a typical ∼10-minute block (**Fig. 2A**). Riding on top of this, pupil size briefly modulated around every decision-interval (**Fig. 2B,C**). On normal trials, time locked to decision-interval onset, we observed a mixture of contrast-related pupil constriction (Barbur et al., 1992) and decision-related pupil dilation (de Gee et al., 2014, 2017, 2020) (**Fig. 2B,C**, top), resulting in overall constriction with respect to pre-trial baseline. We also observed a brief pupil dilation related to the button press (de Gee et al., 2014)) (**Fig. 2B,C**, bottom). Crucially, on the accessory stimulus vs normal trials, pupil-linked arousal was boosted throughout the whole decision-interval before returning to a common baseline (**Fig. 2D**).

**Figure 2.**
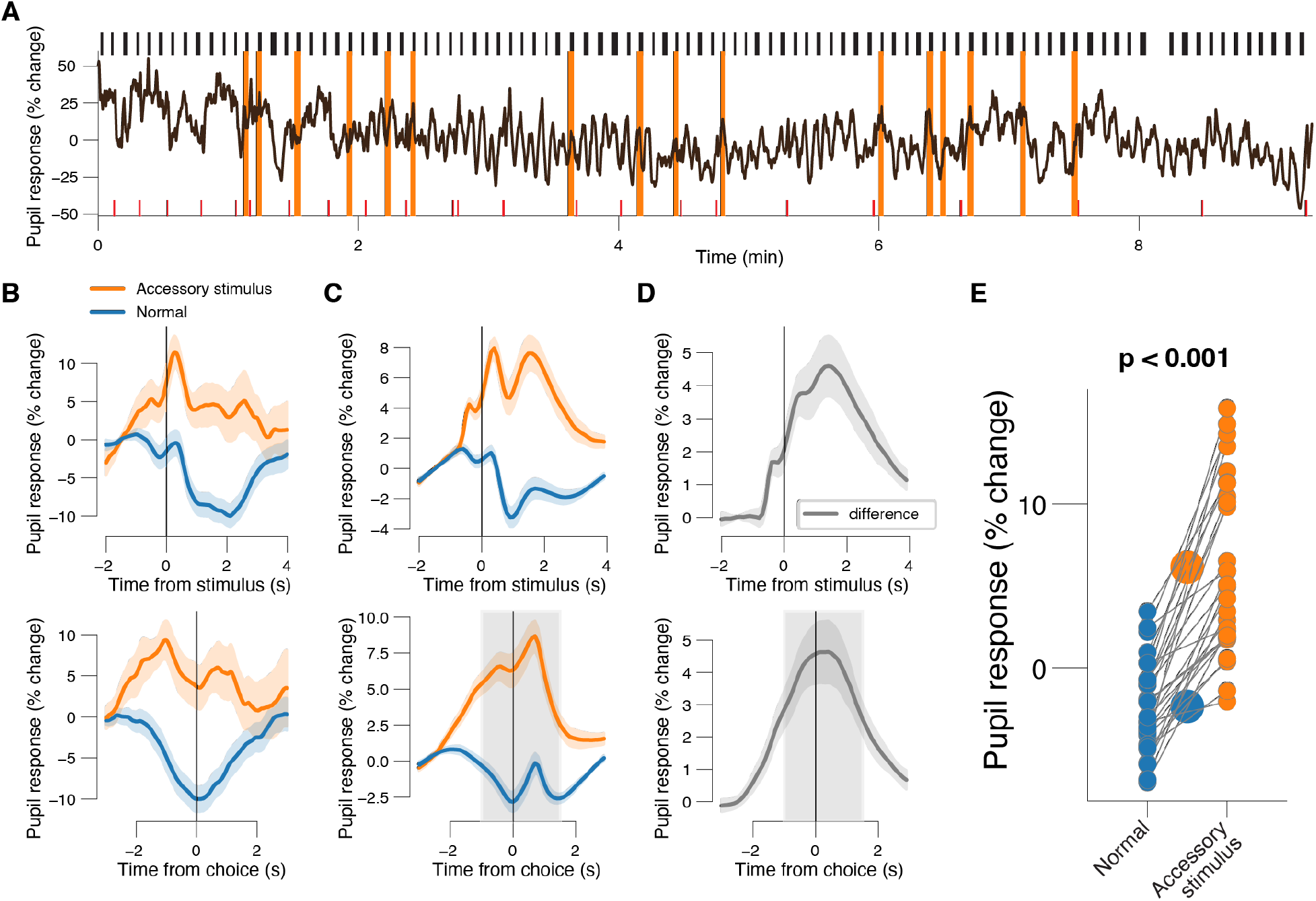
Behavioral task and task-evoked pupil responses. **(A)** Pupil time series from an example block. Black windows (on top), decision intervals; orange windows, accessory stimuli; red windows (below), blinks. **(B)** From the same example block as in A: time course of pupil diameter aligned to decision interval onset (top) or decision interval offset (bottom), and separately for normal and accessory stimulus trials. Shading, 68% confidence intervals across trials (n=80 and n=16 for normal and accessory stimulus trials respectively). **(C)** As B, but for the group average. Shading, 68% confidence interval across subjects (N=29); grey window, interval for computing trial-wise pupil response measures. **(D)** As B, but for the difference between accessory stimulus and normal trials; grey window, interval for computing trial-wise pupil response measures. **(E)** Pupil response amplitude, separately for normal and accessory stimulus trials. Every connecting line is a subject; large data points in the middle are the group averages. Error bars, 68% confidence interval across subjects (N=29); stats, paired sampled t-test.

As before (de Gee et al., 2017, 2020), we computed trial-wise scalar response amplitudes by taking the mean pupil size in the window -1 s from report to 1.5 s after, and subtracting the pre-trial baseline pupil size (Methods). For all subjects, these pupil response amplitudes were higher on the accessory stimulus vs normal trials (**Fig. 2E**). In sum, the accessory stimulus recruited pupil-linked arousal throughout protracted decision-making.

### Accessory stimulus reduces choice bias

We next sought to test the hypothesis that experimentally manipulated phasic pupil-linked arousal reduces the impact of biases on choice behavior. On the normal trials, most subjects (27 out of 29) exhibited a conservative bias (signal detection theoretic criterion (Green & Swets, 1966); Methods), a tendency to choose “no” irrespective of the objective sensory evidence (see x-axis in **Fig. 3A**). We previously showed that phasic arousal predicts a reduction of choice biases irrespective of direction (conservative or liberal) (de Gee et al., 2020). In line with this, we observed a robust relationship between subject’s choice bias on normal trials, and the shift therein caused by the accessory stimulus: subjects with the strongest liberal or conservative biases, exhibited the strongest shift towards a neutral bias (**Fig. 3A**). Correspondingly, the accessory stimulus caused a significant reduction of the absolute value of the bias, measuring the magnitude of bias irrespective of sign (**Fig 3B**).

**Figure 3.**
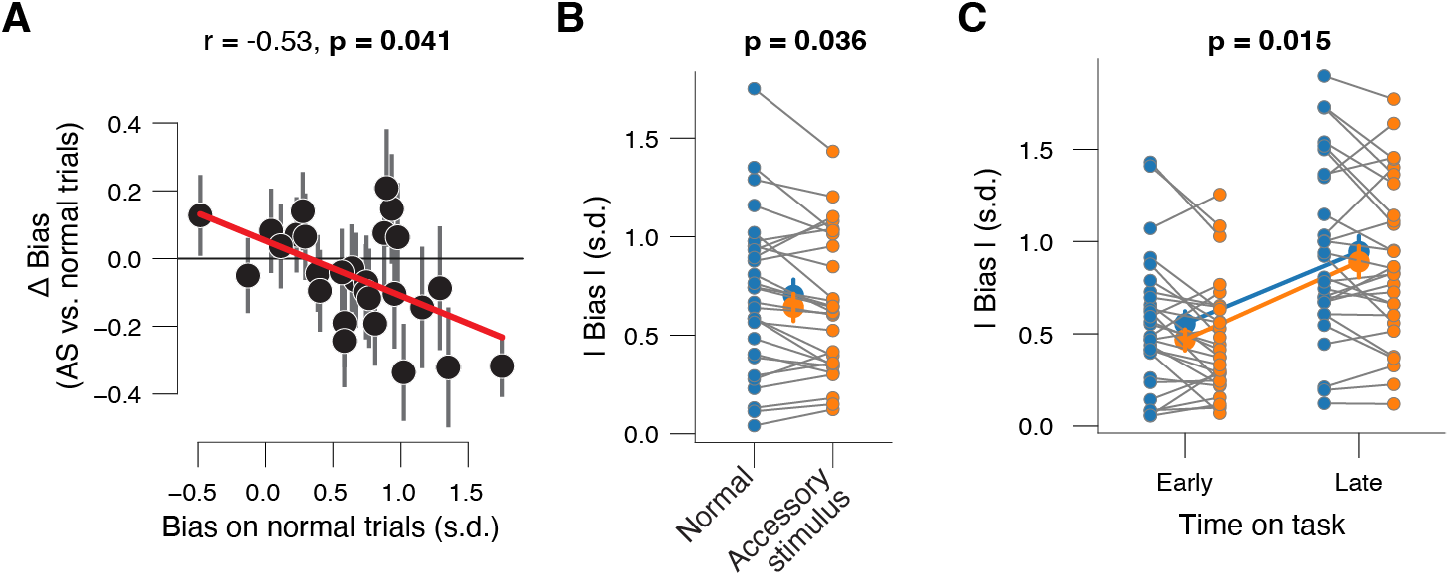
Accessory stimulus reduces choice bias. **(A)** Individual shift in choice bias (signal detection theoretic criterion; Methods) caused by the accessory stimulus, plotted against individual’s choice bias on the normal trials. Data points are individual subjects. Correlation was assessed statistically by Pearson’s correlation coefficient (corrected for reversion to the mean; Methods). Error bars, 68% confidence intervals across bootstraps (n=5K). **(B)** Absolute choice bias, separately for normal and accessory stimulus trials. Every connecting line is a subject; large data points in the middle are the group averages. Error bars, 68% confidence interval across subjects (N=29); stats, paired sampled t-test. **(C)** As B, but separately for normal and accessory stimulus trials and for time on task (Methods). Stats, main effect of accessory stimulus in two-way repeated measures ANOVA (see main text; Methods).

We observed substantial time-on-task effects: subjects tended to get progressively slower, less accurate, and more conservative throughout each block (**Fig. S1B-E**). We performed a two-way repeated measures ANOVA to compare the effects of the accessory stimulus and time on task on absolute bias (**Fig. 3C**; Methods). There was a main effect of the accessory stimulus (F_1,27_ = 6.8, p = 0.015), a main effect of time on task (F_1,27_ = 35.4, p < 0.001), but no interaction (F_1,27_ = 0.4, p = 0.542). Thus, the effect of the accessory stimulus on absolute bias was robust and not dependent on time on task.

In sum, confirming our hypothesis, the accessory stimulus caused a reduction of choice bias.

### Accessory stimulus effect on choice bias is not due to a reduction of response caution

One potential confound in our design was that subjects could control the offset of the slightly aversive auditory white noise, by making a choice (with a button press). This may have caused them to favor speed over sensitivity on the accessory stimulus trials. In line with this, on the accessory stimulus vs normal trials, reaction times were shorter (**Fig. 1B**) and signal detection theoretic sensitivity (d’; Methods; (Green & Swets, 1966)) was reduced (**Fig. S2**). With drift diffusion modeling (**Fig. 4A**; (Bogacz et al., 2006; Ratcliff & McKoon, 2008)) we verified that such a response caution effect did not account for the effect of the accessory stimulus on choice bias.

**Figure 4.**
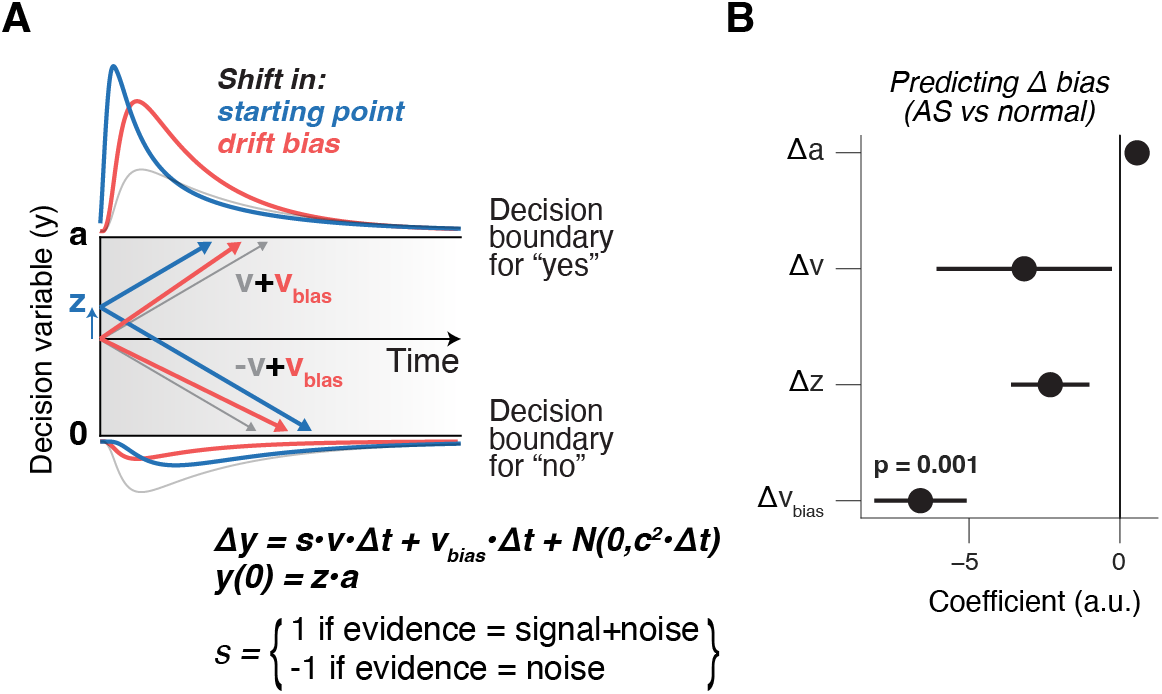
Accessory stimulus reduces choice bias. **(A)** Schematic of drift diffusion model accounting for choices, and their associated RTs. In the equation, *v* is the drift rate. Red and blue curves on the left show expected RT distributions under shifts in either the “starting point” (z; blue) or “drift bias” (v_bias_; red). **(B)** Coefficients from across-subjects multiple linear regression of the change in signal detection theoretic bias (accessory stimulus vs normal trials) on the change in boundary separation (a), change in drift rate (v), change in starting point bias (z) and change in drift bias (v_bias_). Error bars, standard error; stats are FDR-corrected (Methods). See **Fig. S3G-J** for separate scatterplots of partial correlations.

Mimicking our previous approach (de Gee et al., 2017, 2020) we fitted all five main parameters separately for the normal and accessory stimulus trials: 1) boundary separation (controlling response caution); 2) the mean drift rate (overall efficiency of evidence accumulation); 3) non-decision time (the speed of pre-decisional evidence encoding and post-decisional translation of choice into motor response); 4) the starting point of the decision; 5) an evidence-independent bias in the drift (henceforth called “drift bias”). Starting point bias and drift bias can produce the same bias in choice fractions but have distinct effects on the shape of reaction time distributions (**Fig. 4A**; (de Gee et al., 2017, 2020; Urai et al., 2019)): a starting point bias primarily affects relatively fast choices, whereas drift biases accumulate over time and predict biased choices also when reaction times are slower.

The model accounted well for the overall behavior in each task (**Fig. S3A-D**). As expected, the accessory stimulus caused a significant reduction in the boundary separation (**Fig. S3E**). We further observed a trend towards an increased drift rate (**Fig. S3F**) and a reduction of non-decision time (**Fig. S3G**).

Smaller boundary separation on accessory stimulus vs normal trials, when combined with a static drift bias, could in principle account smaller behavioral biases: the drift bias accumulates over time, and cutting the decision process short will reduce its effect. Our analyses deem this scenario unlikely. Across subjects we regressed the shift in bias caused by the accessory stimulus on the shift in all parameters of the drift diffusion model (except non-decision time, Methods). We found that only changes in drift bias (and not changes in boundary separation or any other parameter) predicted the reduction in behavioral bias caused by the accessory stimulus (**Fig. 3B**). This result is in line with our earlier finding that a pupil-predicted reduction of choice bias was due to a change in drift bias, and not in starting point bias (de Gee et al., 2017, 2020),

Taken together, computational modeling converged on the conclusion that changes in drift bias, but not a bias in the starting point of evidence accumulation, implement the pupil-linked shifts in choice bias.

## Discussion

Recent work indicates that pupil-linked phasic arousal signals reduce the impact of prior expectations and biases on decision formation (de Gee et al., 2014, 2017, 2020; Filipowicz et al., 2020; Krishnamurthy et al., 2017; Lewandowska et al., 2019; Murphy et al., 2021; Nassar et al., 2012; Schriver et al., 2020; Urai et al., 2017). Here, using an auditory accessory stimulus, we found that this relationship is causal: on accessory stimulus vs normal trials, pupil-linked phasic arousal was robustly elevated and choice bias was reduced. With computational modeling of behavior, we established that the bias reduction was not due to a change in response caution (i.e., speed-accuracy tradeoff), but due to a change in a bias in the accumulation of evidence leading up to a choice.

We here used pupil responses as a peripheral readout of changes in cortical arousal state (Joshi & Gold, 2020; Larsen & Waters, 2018; McGinley et al., 2015). Changes in pupil diameter have been associated with the activity of the noradrenergic locus coeruleus and the cholinergic basal forebrain in humans (de Gee et al., 2017; Murphy et al., 2014), monkeys (Joshi et al., 2016; Varazzani et al., 2015), and mice (Breton-Provencher & Sur, 2019; Liu et al., 2017; Mridha et al., 2021; Reimer et al., 2016). Recently, pupil size has also been shown to track serotonin and orexin (Cazettes et al., 2021; Grujic et al., 2022). Thus, the exact neuroanatomical and neurochemical source(s) of our observed effects of experimentally controlled phasic arousal on decision-making remain to be determined. Future work should track the activity of multiple neuromodulatory nuclei at once, with brainstem fMRI in humans (Colizoli et al., 2022; de Gee et al., 2017; Mazancieux et al., 2022) or multi-fiber photometry / multi-color 2-photon imaging (of axons) in mice (Mridha et al., 2021; Sych et al., 2019).

Our finding that pupil-linked phasic arousal causally reduces choice bias has important implications for theories about the functional role of these systems in decision-making. First, our results are in line with the idea that (at least some of) the ascending arousal systems that are read out by pupil size are active early enough during a protracted decision process to shape the unfolding decision computations and the subsequent choice outcome (Cheadle et al., 2014; de Gee et al., 2014, 2017). Second, our results are consistent with the idea that neuromodulatory responses alter the balance between “bottom-up” and “top-down” signaling across the cortical hierarchy: cortical sensory regions encode likelihood signals and send these (bottom-up) to association cortex, while participants’ prior beliefs (about target probability, for example) are sent back (top-down) to the lower levels of the hierarchy (Beck et al., 2012; Pouget et al., 2013); neuromodulators might reduce the weight of this prior data in the inference process (Friston, 2010; Hasselmo, 2006; Hsieh et al., 2000; Kimura et al., 1999; Kobayashi et al., 2000; Moran et al., 2013), thereby reducing choice biases.

The auditory accessory stimulus likely recruited the ascending arousal system for at least two reasons. First, the accessory stimulus occurred on a random one-sixth of trials and was therefore surprising. Indeed, pupil-linked phasic arousal is reliably elevated after surprising events (de Gee et al., 2021; Filipowicz et al., 2020; Joshi et al., 2016; Kloosterman et al., 2015; Murphy et al., 2014; Preuschoff & Hart, 2011). Second, the auditory white noise (presented at 70 dB) might be intrinsically arousing, irrespective of the inability of subjects to predict when the sound will be presented. Consistent with this idea, continuously presented background auditory white noise also affects human task performance (Corcoran, 1962; Han et al., 2021; Söderlund et al., 2010). Future research should investigate if the bias-suppressing effect of the accessory stimulus also occurs during more frequent accessory stimuli, for example on 25% of 50% of trials (Lippert et al., 2007; Tona et al., 2016).

One potential confound in our design was that subjects could control the offset of the slightly aversive auditory white noise, by making a choice (with a button press). This may have caused them to favor speed over sensitivity on the accessory stimulus trials. In line with this, computational modeling of behavior indicated that subject’s response caution was indeed lower on the accessory stimulus trials. Crucially, we verified that this did not explain the bias-suppressing effect of the accessory stimulus. Interestingly, the finding that the accessory stimulus did not affect accuracy indicates that its detrimental effect on sensitivity (due to reduced response caution) and its beneficial effect on choice bias cancelled each other out. A reduction of bias should indeed increase accuracy: in our contrast detection task, with signals present on 50% of trials, any bias that deviates from 0 is reduces accuracy.

It is important to establish approaches to reliably manipulate pupil-linked arousal, because (i) it permits the expression of causal questions about the role of neuromodulatory systems in for example perception, learning and decision-making, and (ii) it could lead to practical applications in health (as a cognitive enhancer) and disease (i.e., ADHD). Other non-invasive strategies are pharmacology and transcutaneous vagus nerve stimulation. For example, the noradrenaline re-uptake inhibitor atomoxetine boost pupil-linked arousal (Pfeffer et al., 2021) and has successfully been used in fundamental cognitive neuroscience (Loughnane et al., 2019; Pfeffer et al., 2021; Warren et al., 2016). However, one important drawback of pharmacology is the lack of temporal precision. Another promising approach is vagus nerve stimulation (Mridha et al., 2021; Sharon et al., 2021). However, the optimal parameters for transcutaneous stimulation are currently unknown (Keute et al., 2019). Here we confirm an auditory accessory stimulus as a low-tech approach to precisely control phasic pupil-linked arousal and behavior.

## Methods

### Subjects

Thirty-one healthy subjects (24 females; age range, 18-29 y) participated in the experiment. All subjects had normal or corrected-to-normal vision and gave written informed consent. Subjects received research credit for their participation. The ethics committee of the Psychology Department of the University of Amsterdam approved the experiments. Three subjects were excluded from the analyses: two subjects blinked during more than 20% of decision intervals, and one subject’s accuracy on the normal trials was less than 60% correct.

### Behavioral task

Each trial consisted of three consecutive intervals: (i) the baseline interval (1 s); (ii) the decision interval (terminated by the subject’s response, or after a maximum duration of 3 s); (iii) the inter-trial interval (ITI; uniformly distributed between 3 and 4 s). The baseline interval consisted of green fixation dot on an otherwise blank screen. The decision interval consisted of a green fixation dot and a dynamic noise pattern, with a sinusoidal grating superimposed on 50% of trials. The ITI consisted of a dark grey fixation dot on an otherwise blank screen.

The luminance across all pixels of the dynamic noise pattern (refresh rate: 60 Hz) was kept constant. This pedestal noise pattern had 20% contrast and was refreshed on each frame. On one half of trials (“signal+noise” trials), a sinusoidal grating (5 cycles per degree; vertical orientation) was superimposed on the visual noise for the entire decision interval. The other half of trials (“noise” trials) contained no target signal during the decision interval. Signal presence was randomly selected on each trial, under the constraint that it would occur on 50% of the trials within each block of 96 trials. All stimuli were presented in a Gaussian annulus, with an average distance (±SD) to fixation of 4.8 (1.8) degrees.

Subjects were instructed to report the presence or absence of the signal by pressing one of two response buttons with their left or right index finger once they felt sufficiently certain (free response paradigm). The mapping between perceptual choice and button press (e.g., “yes” –> press right key; “no” –> press left key) was counterbalanced across subjects. At the end of each block of 96 trials, subjects were informed about their performance.

In each block of 96 trials, eight noise trials and eight signal+noise trials (total of 16 trials) were randomly selected as accessory stimulus trials. On these trials, auditory white noise (70 dB) was presented from the start of the baseline interval until subjects terminated the decision interval with a button press.

Throughout the main experiment, the contrast of the target signal was fixed at a level that yielded about 75% correct choices. Each subject’s individual threshold contrast was determined before the main experiment, using an adaptive staircase procedure (Quest). Here, we used a two-interval forced choice variant of the contrast detection task (one interval, signal+noise; the other, noise). The corresponding threshold contrasts yielded a mean accuracy of 73.25% correct (±0.98 % s.e.m.) on the normal trials (**Fig. 1B**).

Visual stimuli were presented on a gamma-corrected monitor with a spatial resolution of 1920 × 1200 pixels, run at a vertical refresh rate of 60 Hz. Auditory stimuli were presented using an IMG Stageline MD-5000DR over-ear headphone, suppressing ambient noise.

Subjects performed between 9 and 14 blocks (distributed over two sessions), yielding a total of 864-1344 trials per subject.

### Eye data acquisition

The left eye’s pupil was tracked at 1000 Hz with an average spatial resolution of 15 to 30 min arc, using an EyeLink 1000 Long Range Mount (SR Research, Osgoode, Ontario, Canada). The eye tracker was calibrated once at the start of each block.

### Analysis of task-evoked pupil responses

#### Preprocessing

Periods of blinks and saccades were detected using the manufacturer’s standard algorithms with default settings. The remaining data analyses were performed using custom-made Python software. We applied to each pupil recording (i) linear interpolation of values measured just before and after each identified blink (interpolation time window, from 150 ms before until 150 ms after blink), (ii) band-pass filtering (third-order Butterworth, passband: 0.01–6 Hz), (iii) removal of pupil responses to blinks and to saccades, by first estimating these responses by means of deconvolution and then removing them from the pupil time series by means of multiple linear regression (Knapen et al., 2016), and (iv) conversion to units of modulation (percent signal change) around the mean of the pupil time series from each block.

#### Quantification of task-evoked pupil responses

We computed task-evoked pupil response measures for each trial as the mean of the pupil diameter modulation values in the window -1 s to 1.5 s from choice (same time window as in (de Gee et al., 2014, 2017, 2020)), minus the mean pupil size during the 0.5 s before the baseline interval.

### Analysis and modeling of choice behavior

We analyzed behavior using custom-made Python software. We excluded the first 10 trials of each block, as we observed that subjects spent these becoming engaged in the task (**Fig. S1**). Reaction time (RT) was defined as the time from decision interval onset until the button press.

#### Signal detection theoretic modeling

We computed the SDT-metric d’ (Green & Swets, 1966) as the difference between z-scores of hit- and false-alarm rates. Criterion was computed as the average of the z-scores of hit- and false-alarm rates, multiplied by -1.

#### Drift diffusion modeling

We used the HDDM 0.9.6 package (Fengler et al., 2021; Wiecki et al., 2013) to fit behavioral data. We allowed the following parameters to vary with condition (normal vs accessory stimulus trials): (i) the separation between both bounds (i.e. response caution); (ii) the mean drift rate across trials; (iii) the non-decision time (sum of the latencies for sensory encoding and motor execution of the choice); (iv) starting point (a bias in the starting point of evidence accumulation toward one of the choice bounds); and (v) drift bias (an evidence independent constant added to the drift). We fitted drift rate variability to all data (i.e., regardless of trial type), to maximize the robustness of our fits.

We fitted the data using Markov-chain Monte Carlo sampling as implemented in the HDDM toolbox (Wiecki et al., 2013). Specifically, we specified a linear model that used categorical dummy-coding to estimate a within-subject effect of the accessory stimulus on each of the five main parameters. Fitting the model to RT distributions for the separate responses (termed ‘stimulus coding’ in Wiecki et al., 2013) enabled the estimation of parameters that could have induced biases towards specific choices. Bayesian MCMC generates full posterior distributions over parameter estimates, quantifying not only the most likely parameter value but also the uncertainty associated with that estimate. The hierarchical nature of the model assumes that all observers in a data set are drawn from a group, with specific group-level prior distributions that are informed by the literature. In practice, this results in more stable parameter estimates for individual subjects, who are constrained by the group-level inference. The hierarchical nature of the model also minimizes the risk of overfitting the data (Katahira, 2016; Vandekerckhove et al., 2011; Wiecki et al., 2013). Together, this allowed us to vary all main parameters simultaneously with condition, so that they could “compete” capturing the reduction of choice bias caused by the accessory stimulus).

We ran five separate Markov chains with 12500 samples each. Of those, 2500 were discarded as burn-in. Individual parameter estimates were then estimated from the posterior distributions across the resulting 10000 samples. All group-level chains were visually inspected to ensure convergence. In addition, we computed the Gelman-Rubin R statistic (which compares within-chain and between-chain variance) and checked that all group-level parameters had an R between 0.99 and 1.01.

### Statistical comparisons

We used a paired sample t-test to test for significant differences between behavioral estimates (**Fig. 1 & Fig. 3**) task-evoked pupil responses (**Fig. 2**) from different trial categories. We used a 2 × 2 repeated measures ANOVA to test for the main effect of trial category and time-on-task, and their interaction (**Fig. 3**).

We used Pearson correlation coefficient to quantify the strength of the relationship between subject’s bias on normal trials and the change in bias caused by the accessory stimulus (**Fig 3**). We performed permutation analyses to obtain the distribution of correlation coefficients predicted exclusively by regression to the mean. This was done by computing the above-described across-subject correlation 10K times, each time using randomly assigned “normal” vs “accessory stimulus” labels of each subject. We then compared our observed correlation coefficient (reflecting the combined effects of regression to the mean and the true relationship) against this permutation distribution. This analysis suggested that the observed relationship was stronger than expected based on regression to the mean (proportion of permutations larger than the observed correlation was 0.041).

We used multiple linear regression to regres the shift in bias caused by the accessory stimulus on the shift in all parameters of the drift diffusion model (except non-decision time; **Fig. 4**). We left non-decision time out, because this parameter only shifts the reaction time distribution and cannot change choice bias (also not in combination with other parameters). The resulting p-values (one for each parameter) were corrected for multiple comparisons using the false discovery rate (FDR).

Statistical tests were performed at the group level, using the individual subjects’ mean parameters as observations. All tests were performed two-tailed. All error bars are 68% bootstrap confidence intervals of the mean, unless stated otherwise.

## Data availability

Data will be made publicly available upon publication.

## Code availability

Analysis scripts will be made publicly available upon publication.

## Acknowledgements

We thank Simon van Gaal for helpful comments on the behavioral paradigm.

## Author contributions

Conceptualization: THD, MJM, JWdG

Investigation: BMB, VGK, RdV

Formal analysis: JWdG

Writing—original draft: JWdG

Writing—review and editing: THD, MJM, JWdG

Supervision: JWdG

## Supplementary figures

**Figure S1.**
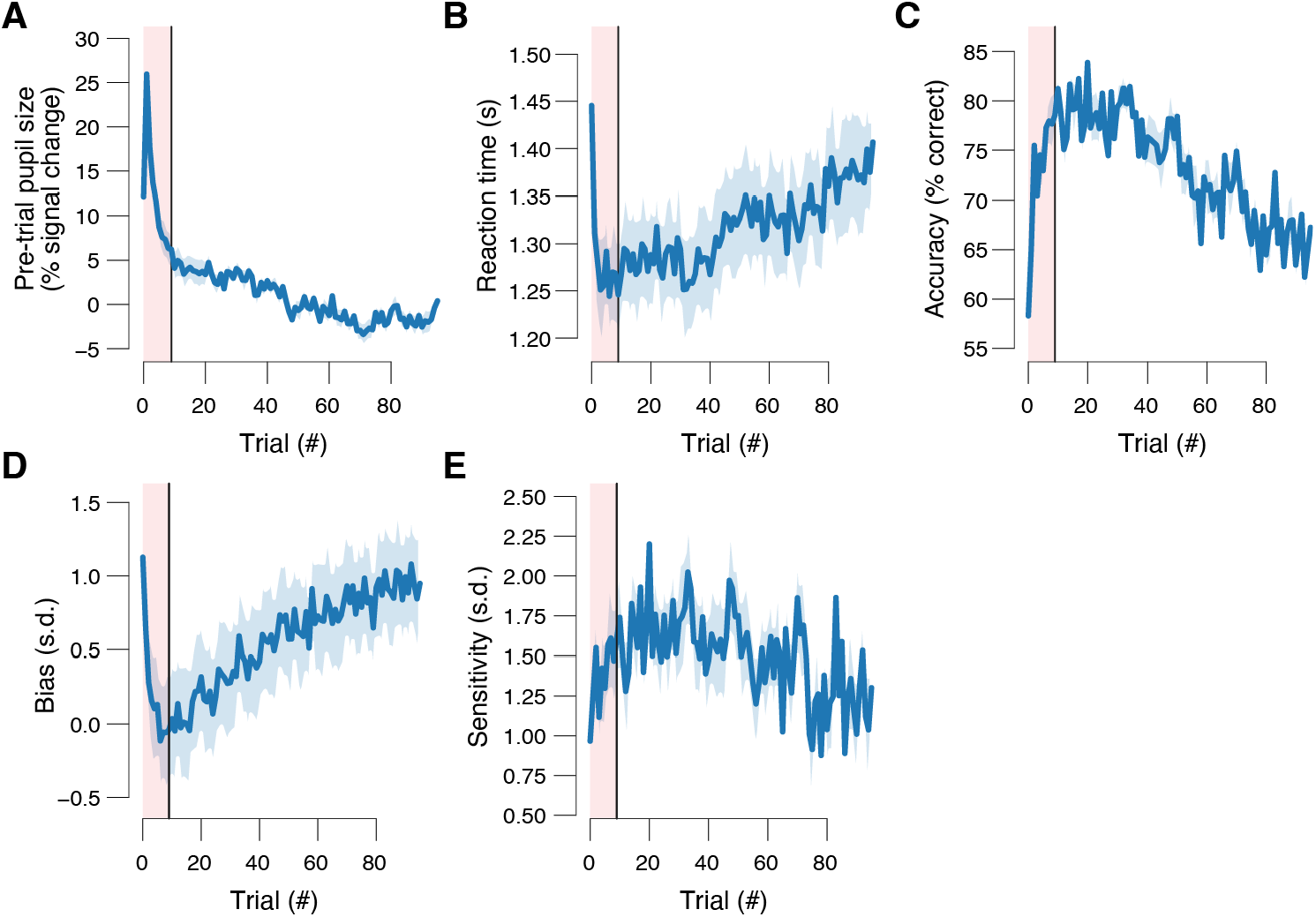
**(A)** Baseline pupil size separately per trial within a block. Shading, 68% confidence interval across blocks (n=334); red window, trials excluded from analyses (Methods). **(B)** As A, but for accuracy. **(C)** As A, but for reaction time. **(D)** Signal detection theoretic sensitivity (Methods) separately per trial within a block. Shading, 68% confidence interval across bootstraps (n=250); red window, trials excluded from analyses (Methods). **(E)** As D, but for signal detection theoretic criterion (Methods).

**Figure S2.**
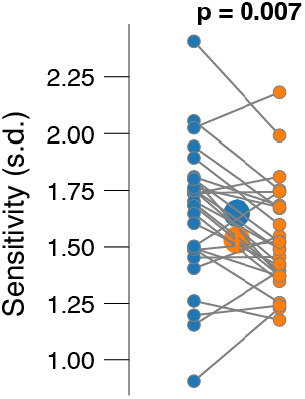
Signal detection theoretic sensitivity (Methods), separately for normal and accessory stimulus trials. Every connecting line is a subject; large data points in the middle are the group averages. Error bars, 68% confidence interval across subjects (N=29); stats, paired sampled t-test.

**Figure S3.**
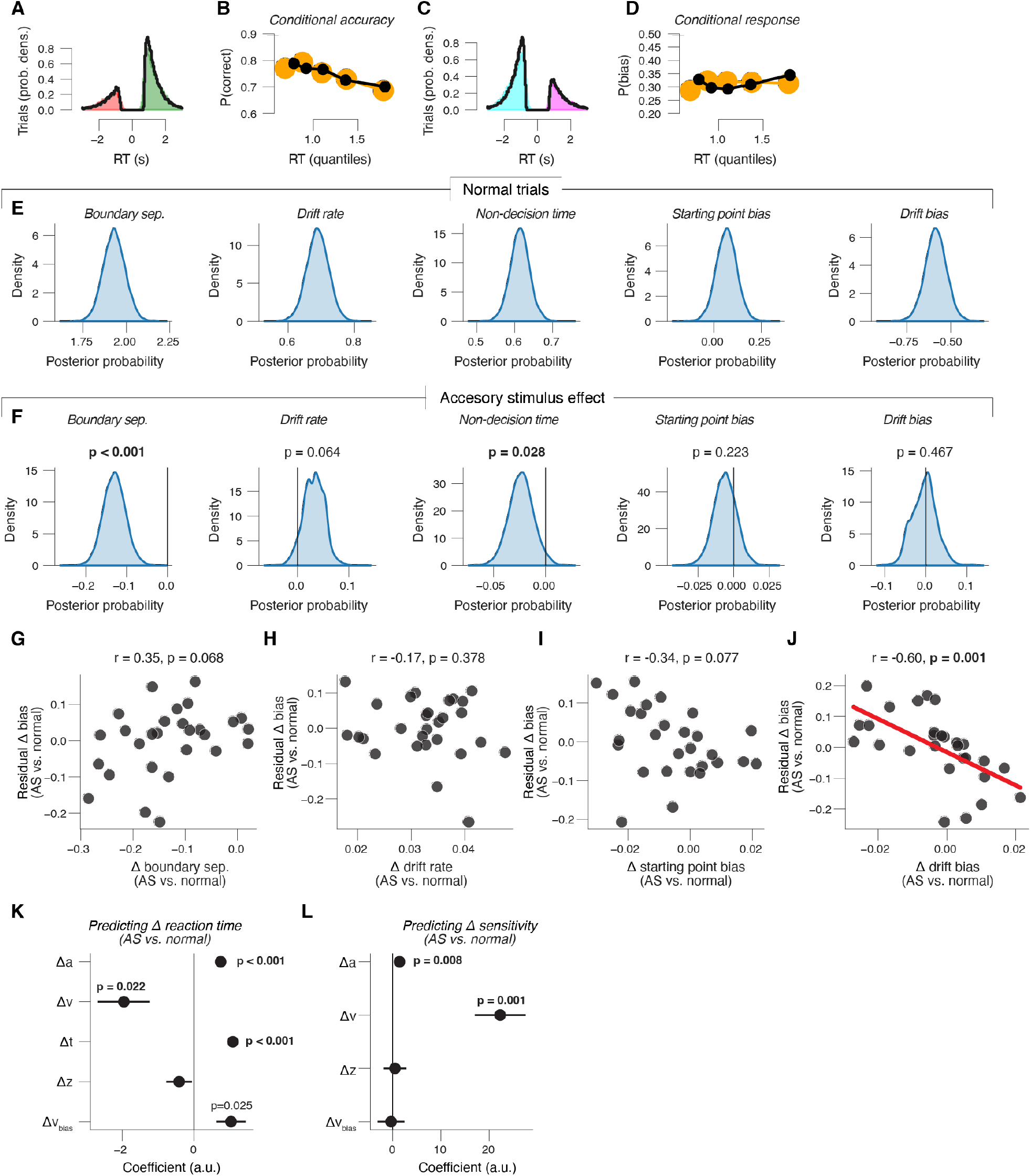
**(A)** Group average RT distributions on normal trials, separately for correct (green) and error (red) choices. Black line, model fit. **(B)** Conditional accuracy functions were generated by dividing RT distributions into five quantiles and computing the fraction of correct choices in each quantile. Black line, model fit. **(C)** As A, but for yes (magenta) and no (cyan) choices. **(D)** Conditional response functions were generated by dividing RT distributions into five quantiles and computing the fraction of yes choices in each quantile. Black line, model fit. **(E)** Group-level posterior probability densities for means of parameters on the normal trials. **(F)** Group-level posterior probability densities for accessory stimulus effect (accessory stimulus vs normal trials) on all parameters. **(G)** Residual change in choice bias (after regressing out effects of change in drift rate, starting point bias and drift bias) plotted against the change in boundary separation. Every data point is a subject. **(H)** Residual change in choice bias (after regressing out effects of change in boundary separation, starting point bias and drift bias) plotted against the change in drift rate. Every data point is a subject. **(I)** Residual change in choice bias (after regressing out effects of change in boundary separation, drift rate and drift bias) plotted against the change in starting point bias. Every data point is a subject. **(J)** Residual change in choice bias (after regressing out effects of change in boundary separation, drift rate and starting point bias) plotted against the change in drift bias. Every data point is a subject. **(K)** Coefficients from across-subjects multiple linear regression of the change in reaction time (accessory stimulus vs normal trials) on the change in boundary separation (a), change in drift rate (v), change in non-decision time (t), change in starting point bias (z) and change in drift bias (v_bias_). Error bars, standard error; stats are FDR-corrected (Methods). **(L)** As K, but for the change in signal detection theoretic sensitivity.

## References

Aston-Jones, G., & Cohen, J. D. (2005). An Integrative Theory of Locus Coeruleus-Norepinephrine Function: Adaptive Gain and Optimal Performance. Annual Review of Neuroscience, 28(1), 403–450. https://doi.org/10.1146/annurev.neuro.28.061604.135709

Barbur, J. L., Harlow, A. J., & Sahraie, A. (1992). Pupillary responses to stimulus structure, colour and movement. Ophthalmic & Physiological Optics: The Journal of the British College of Ophthalmic Opticians (Optometrists), 12(2), 137–141. https://doi.org/10.1111/j.1475-1313.1992.tb00276.x

Beck, J. M., Ma, W. J., Pitkow, X., Latham, P. E., & Pouget, A. (2012). Not noisy, just wrong: The role of suboptimal inference in behavioral variability. Neuron, 74(1), 30–39.

Bernstein, I. H., Clark, M. H., & Edelstein, B. A. (1969). Effects of an auditory signal on visual reaction time. Journal of Experimental Psychology, 80(3, Pt.1), 567–569. https://doi.org/10.1037/h0027444

Bogacz, R., Brown, E., Moehlis, J., Holmes, P., & Cohen, J. D. (2006). The physics of optimal decision making: A formal analysis of models of performance in two-alternative forced-choice tasks. Psychological Review, 113(4), 700–765.

Breton-Provencher, V., Drummond, G. T., Feng, J., Li, Y., & Sur, M. (2022). Spatiotemporal dynamics of noradrenaline during learned behaviour. Nature, 606(7915), 732–738. https://doi.org/10.1038/s41586-022-04782-2

Breton-Provencher, V., & Sur, M. (2019). Active control of arousal by a locus coeruleus GABAergic circuit. Nature Neuroscience, 22(2), 218–228. https://doi.org/10.1038/s41593-018-0305-z

Cazettes, F., Reato, D., Morais, J. P., Renart, A., & Mainen, Z. F. (2021). Phasic Activation of Dorsal Raphe Serotonergic Neurons Increases Pupil Size. Current Biology, 31(1), 192-197.e4. https://doi.org/10.1016/j.cub.2020.09.090

Cheadle, S., Wyart, V., Tsetsos, K., Myers, N., de Gardelle, V., Herce Castañón, S., & Summerfield, C. (2014). Adaptive gain control during human perceptual choice. Neuron, 81(6), 1429–1441.

Colizoli, O., de Gee, J. W., van der Zwaag, W., & Donner, T. H. (2022). Functional magnetic resonance imaging responses during perceptual decision-making at 3 and 7 T in human cortex, striatum, and brainstem. Human Brain Mapping, 43(4), 1265–1279. https://doi.org/10.1002/hbm.25719

Corcoran, D. W. J. (1962). Noise and Loss of Sleep. Quarterly Journal of Experimental Psychology, 14(3), 178–182. https://doi.org/10.1080/17470216208416533

de Gee, J. W., Colizoli, O., Kloosterman, N. A., Knapen, T., Nieuwenhuis, S., & Donner, T. H. (2017). Dynamic modulation of decision biases by brainstem arousal systems. ELife, 6, 309.

de Gee, J. W., Correa, C. M. C., Weaver, M., Donner, T. H., & van Gaal, S. (2021). Pupil Dilation and the Slow Wave ERP Reflect Surprise about Choice Outcome Resulting from Intrinsic Variability in Decision Confidence. Cerebral Cortex, bhab032. https://doi.org/10.1093/cercor/bhab032

de Gee, J. W., Knapen, T., & Donner, T. H. (2014). Decision-related pupil dilation reflects upcoming choice and individual bias. Proceedings of the National Academy of Sciences of the United States of America, 111(5), E618–25.

de Gee, J. W., Tsetsos, K., Schwabe, L., Urai, A. E., McCormick, D., McGinley, M. J., & Donner, T. H. (2020). Pupil-linked phasic arousal predicts a reduction of choice bias across species and decision domains. ELife, 9, e54014. https://doi.org/10.7554/eLife.54014

de Lange, F. P., Rahnev, D. A., Donner, T. H., & Lau, H. (2013). Prestimulus oscillatory activity over motor cortex reflects perceptual expectations. The Journal of Neuroscience, 33(4), 1400–1410.

Fengler, A., Govindarajan, L. N., Chen, T., & Frank, M. J. (2021). Likelihood approximation networks (LANs) for fast inference of simulation models in cognitive neuroscience. ELife, 10, e65074. https://doi.org/10.7554/eLife.65074

Filipowicz, A. L., Glaze, C. M., Kable, J. W., & Gold, J. I. (2020). Pupil diameter encodes the idiosyncratic, cognitive complexity of belief updating. ELife, 9, e57872. https://doi.org/10.7554/eLife.57872

Friston, K. (2010). The free-energy principle: A unified brain theory? Nature Reviews Neuroscience, 11(2), 127–138.

Froemke, R. C. (2015). Plasticity of Cortical Excitatory-Inhibitory Balance. Annual Review of Neuroscience, 38(1), 195–219. https://doi.org/10.1146/annurev-neuro-071714-034002

Gilzenrat, M. S., Nieuwenhuis, S., Jepma, M., & Cohen, J. D. (2010). Pupil diameter tracks changes in control state predicted by the adaptive gain theory of locus coeruleus function. Cognitive, Affective & Behavioral Neuroscience, 10(2), 252–269.

Green, D. M., & Swets, J. A. (1966). Signal detection theory and psychophysics. 1966. New York.

Gritton, H. J., Howe, W. M., Mallory, C. S., Hetrick, V. L., Berke, J. D., & Sarter, M. (2016). Cortical cholinergic signaling controls the detection of cues. Proceedings of the National Academy of Sciences, 113(8), E1089–E1097. https://doi.org/10.1073/pnas.1516134113

Grujic, N., Tesmer, A., Bracey, E. F., Peleg-Raibstein, D., & Burdakov, D. (2022). Control and coding of pupil size by hypothalamic orexin neurons (p. 2022.04.12.488026). bioRxiv. https://doi.org/10.1101/2022.04.12.488026

Hackley, S. A., Langner, R., Rolke, B., Erb, M., Grodd, W., & Ulrich, R. (2009). Separation of phasic arousal and expectancy effects in a speeded reaction time task via fMRI. Psychophysiology, 46(1), 163–171. https://doi.org/10.1111/j.1469-8986.2008.00722.x

Hackley, S. A., & Valle-Inclán, F. (1998). Automatic alerting does not speed late motoric processes in a reaction-time task. Nature, 391(6669), 786–788.

Hackley, S. A., & Valle-Inclán, F. (1999). Accessory stimulus effects on response selection: Does arousal speed decision making? Journal of Cognitive Neuroscience, 11(3), 321–329.

Han, S., Zhu, R., & Ku, Y. (2021). Background white noise and speech facilitate visual working memory. European Journal of Neuroscience, 54(7), 6487–6496. https://doi.org/10.1111/ejn.15455

Hanks, T. D., Mazurek, M. E., Kiani, R., Hopp, E., & Shadlen, M. N. (2011). Elapsed decision time affects the weighting of prior probability in a perceptual decision task. The Journal of Neuroscience : The Official Journal of the Society for Neuroscience, 31(17), 6339–6352. https://doi.org/10.1523/JNEUROSCI.5613-10.2011

Harris, K. D., & Thiele, A. (2011). Cortical state and attention. Nature Reviews Neuroscience, 12(9), 509–523.

Hasselmo, M. E. (2006). The Role of Acetylcholine in Learning and Memory. Current Opinion in Neurobiology, 16(6), 710–715. https://doi.org/10.1016/j.conb.2006.09.002

Hershenson, M. (1962). Reaction time as a measure of intersensory facilitation. Journal of Experimental Psychology, 63(3), 289–293. https://doi.org/10.1037/h0039516

Hsieh, C. Y., Cruikshank, S. J., & Metherate, R. (2000). Differential modulation of auditory thalamocortical and intracortical synaptic transmission by cholinergic agonist. Brain Research, 880(1–2), 51–64.

Jepma, M., Wagenmakers, E.-J., Band, G. P. H., & Nieuwenhuis, S. (2009). The Effects of Accessory Stimuli on Information Processing: Evidence from Electrophysiology and a Diffusion Model Analysis. Journal of Cognitive Neuroscience, 21(5), 847–864. https://doi.org/10.1162/jocn.2009.21063

Joshi, S., & Gold, J. I. (2020). Pupil Size as a Window on Neural Substrates of Cognition. Trends in Cognitive Sciences. https://doi.org/10.1016/j.tics.2020.03.005

Joshi, S., & Gold, J. I. (2022). Context-dependent relationships between locus coeruleus firing patterns and coordinated neural activity in the anterior cingulate cortex. ELife, 11, e63490. https://doi.org/10.7554/eLife.63490

Joshi, S., Li, Y., Kalwani, R. M., & Gold, J. I. (2016). Relationships between Pupil Diameter and Neuronal Activity in the Locus Coeruleus, Colliculi, and Cingulate Cortex. Neuron, 89(1), 221–234.

Jun, E. J., Bautista, A. R., Nunez, M. D., Allen, D. C., Tak, J. H., Alvarez, E., & Basso, M. A. (2021). Causal role for the primate superior colliculus in the computation of evidence for perceptual decisions. Nature Neuroscience, 24(8), 1121–1131. https://doi.org/10.1038/s41593-021-00878-6

Katahira, K. (2016). How hierarchical models improve point estimates of model parameters at the individual level. Journal of Mathematical Psychology, 73, 37–58. https://doi.org/10.1016/j.jmp.2016.03.007

Keute, M., Demirezen, M., Graf, A., Mueller, N. G., & Zaehle, T. (2019). No modulation of pupil size and event-related pupil response by transcutaneous auricular vagus nerve stimulation (taVNS). Scientific Reports, 9(1), 1–10. https://doi.org/10.1038/s41598-019-47961-4

Kimura, F., Fukuda, M., & Tsumoto, T. (1999). Acetylcholine suppresses the spread of excitation in the visual cortex revealed by optical recording: Possible differential effect depending on the source of input. The European Journal of Neuroscience, 11(10), 3597–3609.

Kloosterman, N. A., de Gee, J. W., Werkle-Bergner, M., Lindenberger, U., Garrett, D. D., & Fahrenfort, J. J. (2019). Humans strategically shift decision bias by flexibly adjusting sensory evidence accumulation. ELife, 8, e37321. https://doi.org/10.7554/eLife.37321

Kloosterman, N. A., Meindertsma, T., Loon, A. M., Lamme, V. A. F., Bonneh, Y. S., & Donner, T. H. (2015). Pupil size tracks perceptual content and surprise. European Journal of Neuroscience, 41(8), 1068–1078.

Knapen, T., de Gee, J. W., Brascamp, J., Nuiten, S., Hoppenbrouwers, S., & Theeuwes, J. (2016). Cognitive and Ocular Factors Jointly Determine Pupil Responses under Equiluminance. PLOS ONE, 11(5), e0155574.

Kobayashi, M., Imamura, K., Sugai, T., Onoda, N., Yamamoto, M., Komai, S., & Watanabe, Y. (2000). Selective suppression of horizontal propagation in rat visual cortex by norepinephrine. The European Journal of Neuroscience, 12(1), 264–272.

Krishnamurthy, K., Nassar, M. R., Sarode, S., & Gold, J. I. (2017). Arousal-related adjustments of perceptual biases optimize perception in dynamic environments. Nature Human Behaviour, 1, 0107.

Larsen, R. S., & Waters, J. (2018). Neuromodulatory Correlates of Pupil Dilation. Frontiers in Neural Circuits, 12.

Lee, S.-H., & Dan, Y. (2012). Neuromodulation of brain states. Neuron, 76(1), 209–222.

Lewandowska, K., Gągol, A., Sikora-Wachowicz, B., Marek, T., & Fąfrowicz, M. (2019). Saying “yes” when you want to say “no”—Pupil dilation reflects evidence accumulation in a visual working memory recognition task. International Journal of Psychophysiology, 139, 18–32. https://doi.org/10.1016/j.ijpsycho.2019.03.001

Lippert, M., Logothetis, N. K., & Kayser, C. (2007). Improvement of visual contrast detection by a simultaneous sound. Brain Research, 1173, 102–109. https://doi.org/10.1016/j.brainres.2007.07.050

Liu, Y., Rodenkirch, C., Moskowitz, N., Schriver, B., & Wang, Q. (2017). Dynamic Lateralization of Pupil Dilation Evoked by Locus Coeruleus Activation Results from Sympathetic, Not Parasympathetic, Contributions. Cell Reports, 20(13), 3099–3112.

Loughnane, G. M., Brosnan, M. B., Barnes, J. J. M., Dean, A., Nandam, S. L., O’Connell, R. G., & Bellgrove, M. A. (2019). Catecholamine Modulation of Evidence Accumulation during Perceptual Decision Formation: A Randomized Trial. Journal of Cognitive Neuroscience, 31(7), 1044–1053. https://doi.org/10.1162/jocn_a_01393

Lovelace, C. T., Stein, B. E., & Wallace, M. T. (2003). An irrelevant light enhances auditory detection in humans: A psychophysical analysis of multisensory integration in stimulus detection. Cognitive Brain Research, 17(2), 447–453. https://doi.org/10.1016/S0926-6410(03)00160-5

Mazancieux, A., Mauconduit, F., Amadon, A., Gee, J. W. de, Donner, T., & Meyniel, F. (2022). Brainstem fMRI signaling of surprise across different types of deviant stimuli (p. 2022.07.25.501374). bioRxiv. https://doi.org/10.1101/2022.07.25.501374

McCormick, D. A., Nestvogel, D. B., & He, B. J. (2020). Neuromodulation of Brain State and Behavior. Annual Review of Neuroscience, 43, 391–415. https://doi.org/10.1146/annurev-neuro-100219-105424

McGinley, M. J., Vinck, M., Reimer, J., Batista-Brito, R., Zagha, E., Cadwell, C. R., Tolias, A. S., Cardin, J. A., & McCormick, D. A. (2015). Waking State: Rapid Variations Modulate Neural and Behavioral Responses. Neuron, 87(6), 1143–1161.

Miller, J., Franz, V., & Ulrich, R. (1999). Effects of auditory stimulus intensity on response force in simple, go/no-go, and choice RT tasks. Perception & Psychophysics, 61(1), 107–119. https://doi.org/10.3758/BF03211952

Mochol, G., Kiani, R., & Moreno-Bote, R. (2021). Prefrontal cortex represents heuristics that shape choice bias and its integration into future behavior. Current Biology, 31(6), 1234-1244.e6. https://doi.org/10.1016/j.cub.2021.01.068

Moran, R. J., Campo, P., Symmonds, M., Stephan, K. E., Dolan, R. J., & Friston, K. J. (2013). Free energy, precision and learning: The role of cholinergic neuromodulation. The Journal of Neuroscience, 33(19), 8227–8236.

Mridha, Z., de Gee, J. W., Shi, Y., Alkashgari, R., Williams, J., Suminski, A., Ward, M. P., Zhang, W., & McGinley, M. J. (2021). Graded recruitment of pupil-linked neuromodulation by parametric stimulation of the vagus nerve. Nature Communications, 12(1), 1539. https://doi.org/10.1038/s41467-021-21730-2

Murphy, P. R., O’Connell, R. G., O’Sullivan, M., Robertson, I. H., & Balsters, J. H. (2014). Pupil diameter covaries with BOLD activity in human locus coeruleus. Human Brain Mapping, 35(8), 4140–4154.

Murphy, P. R., Wilming, N., Hernandez-Bocanegra, D. C., Prat-Ortega, G., & Donner, T. H. (2021). Adaptive circuit dynamics across human cortex during evidence accumulation in changing environments. Nature Neuroscience, 24(7), 987–997. https://doi.org/10.1038/s41593-021-00839-z

Nassar, M. R., Rumsey, K. M., Wilson, R. C., Parikh, K., Heasly, B., & Gold, J. I. (2012). Rational regulation of learning dynamics by pupil-linked arousal systems. Nature Neuroscience, 15(7), 1040–1046.

Nickerson, R. S. (1973). Intersensory facilitation of reaction time: Energy summation or preparation enhancement? Psychological Review, 80(6), 489–509. https://doi.org/10.1037/h0035437

Petersen, A., Petersen, A. H., Bundesen, C., Vangkilde, S., & Habekost, T. (2017). The effect of phasic auditory alerting on visual perception. Cognition, 165, 73–81. https://doi.org/10.1016/j.cognition.2017.04.004

Pfeffer, T., Ponce-Alvarez, A., Tsetsos, K., Meindertsma, T., Gahnström, C. J. Brink, R. L. van den, Nolte, G., Engel, A. K., Deco, G., & Donner, T. H. (2021). Circuit mechanisms for the chemical modulation of cortex-wide network interactions and behavioral variability. Science Advances, 7(29), eabf5620. https://doi.org/10.1126/sciadv.abf5620

Pouget, A., Beck, J. M., Ma, W. J., & Latham, P. E. (2013). Probabilistic brains: Knowns and unknowns. Nature Neuroscience, 16(9), 1170–1178.

Preuschoff, K., & Hart, B. M. (2011). Pupil dilation signals surprise: Evidence for noradrenaline’s role in decision making. Frontiers in \ldots.

Ratcliff, R., & McKoon, G. (2008). The diffusion decision model: Theory and data for two-choice decision tasks. Neural Computation, 20(4), 873–922.

Reimer, J., McGinley, M. J., Liu, Y., Rodenkirch, C., Wang, Q., McCormick, D. A., & Tolias, A. S. (2016). Pupil fluctuations track rapid changes in adrenergic and cholinergic activity in cortex. Nature Communications, 7, 13289.

Schriver, B. J., Perkins, S. M., Sajda, P., & Wang, Q. (2020). Interplay between components of pupil-linked phasic arousal and its role in driving behavioral choice in Go/No-Go perceptual decision-making. Psychophysiology, 00, e13565. https://doi.org/10.1111/psyp.13565

Shadlen, M. N., & Kiani, R. (2013). Decision making as a window on cognition. Neuron, 80(3), 791–806.

Sharon, O., Fahoum, F., & Nir, Y. (2021). Transcutaneous Vagus Nerve Stimulation in Humans Induces Pupil Dilation and Attenuates Alpha Oscillations. Journal of Neuroscience, 41(2), 320–330. https://doi.org/10.1523/JNEUROSCI.1361-20.2020

Siegel, M., Engel, A. K., & Donner, T. H. (2011). Cortical network dynamics of perceptual decision-making in the human brain. Frontiers in Human Neuroscience, 5, 21.

Söderlund, G. B., Sikström, S., Loftesnes, J. M., & Sonuga-Barke, E. J. (2010). The effects of background white noise on memory performance in inattentive school children. Behavioral and Brain Functions, 6(1), 55. https://doi.org/10.1186/1744-9081-6-55

Stahl, J., & Rammsayer, T. H. (2005). Accessory stimulation in the time course of visuomotor information processing: Stimulus intensity effects on reaction time and response force. Acta Psychologica, 120(1), 1–18. https://doi.org/10.1016/j.actpsy.2005.02.003

Steinmetz, N. A., Zatka-Haas, P., Carandini, M., & Harris, K. D. (2019). Distributed coding of choice, action and engagement across the mouse brain. Nature, 576(7786), 266–273. https://doi.org/10.1038/s41586-019-1787-x

Stoffels, E. J., Van Der Molen, M. W., & Keuss, P. J. G. (1985). Intersensory facilitation and inhibition: Immediate arousal and location effects of auditory noise on visual choice reaction time. Acta Psychologica, 58(1), 45–62. https://doi.org/10.1016/0001-6918(85)90033-2

Summerfield, C., & de Lange, F. P. (2014). Expectation in perceptual decision making: Neural and computational mechanisms. Nature Reviews Neuroscience, 15(11), 745–756.

Sych, Y., Chernysheva, M., Sumanovski, L. T., & Helmchen, F. (2019). High-density multi-fiber photometry for studying large-scale brain circuit dynamics. Nature Methods, 16(6), 553–560. https://doi.org/10.1038/s41592-019-0400-4

Tona, K.-D., Murphy, Peter. R., Brown, S. B. R. E., & Nieuwenhuis, S. (2016). The accessory stimulus effect is mediated by phasic arousal: A pupillometry study. Psychophysiology, 53(7), 1108–1113. https://doi.org/10.1111/psyp.12653

Urai, A. E., Braun, A., & Donner, T. H. (2017). Pupil-linked arousal is driven by decision uncertainty and alters serial choice bias. Nature Communications, 8, 14637.

Urai, A. E., de Gee, J. W., Tsetsos, K., & Donner, T. H. (2019). Choice history biases subsequent evidence accumulation. ELife, 8, e46331. https://doi.org/10.7554/eLife.46331

van Vugt, B., Dagnino, B., Vartak, D., Safaai, H., Panzeri, S., Dehaene, S., & Roelfsema, P. R. (2018). The threshold for conscious report: Signal loss and response bias in visual and frontal cortex. Science, 360(6388), 537–542. https://doi.org/10.1126/science.aar7186

Vandekerckhove, J., Tuerlinckx, F., & Lee, M. D. (2011). Hierarchical diffusion models for two-choice response times. Psychological Methods, 16(1), 44–62. https://doi.org/10.1037/a0021765

Varazzani, C., San-Galli, A., Gilardeau, S., & Bouret, S. (2015). Noradrenaline and Dopamine Neurons in the Reward/Effort Trade-Off: A Direct Electrophysiological Comparison in Behaving Monkeys. Journal of Neuroscience, 35(20), 7866–7877.

Wang, X.-J. (2008). Decision making in recurrent neuronal circuits. Neuron, 60(2), 215–234.

Warren, C. M., Eldar, E., van den Brink, R. L., Tona, K.-D., van der Wee, N. J., Giltay, E. J., van Noorden, M. S., Bosch, J. A., Wilson, R. C., Cohen, J. D., & Nieuwenhuis, S. (2016). Catecholamine-Mediated Increases in Gain Enhance the Precision of Cortical Representations. The Journal of Neuroscience, 36(21), 5699–5708.

Wiecki, T. V., Sofer, I., & Frank, M. J. (2013). HDDM: Hierarchical Bayesian estimation of the Drift-Diffusion Model in Python. Frontiers in Neuroinformatics, 7, 14.

